# Alternative Complement Pathway Regulates Thermogenic Fat Function via Adipsin and Adipocyte-derived C3aR1 in a Sex-dependent Fashion

**DOI:** 10.1101/2022.12.30.522320

**Authors:** Lunkun Ma, Ankit Gilani, Alfonso Rubio-Navarro, Eric Cortada-Almar, Ang Li, Shannon M. Reilly, Liling Tang, James C. Lo

## Abstract

Thermogenesis in beige/brown adipose tissues can be leveraged to combat metabolic disorders such as type 2 diabetes and obesity. The complement system plays pleiotropic roles in metabolic homeostasis and organismal energy balance with canonical effects on immune cells and non-canonical effects on non-immune cells. The adipsin/C3a/C3aR1 pathway stimulates insulin secretion and sustains pancreatic beta cell mass. However, its role in adipose thermogenesis has not been defined. Here, we show that *Adipsin* knockout mice exhibit increased energy expenditure and white adipose tissue (WAT) browning. C3a, a downstream product of adipsin, is generated from complement component 3 and decreases *Ucp1* expression in subcutaneous adipocytes. In addition, adipocyte-specific *C3aR1* knockout male mice show enhanced WAT thermogenesis and increased respiration. In stark contrast, adipocyte-specific *C3aR1* knockout female mice display decreased brown fat thermogenesis and are cold intolerant. Female mice express lower levels of *Adipsin* in thermogenic adipocytes and adipose tissues than males. *C3aR1* is also lower in female subcutaneous adipose tissue than males. Collectively, these results reveal sexual dimorphism in the adipsin/C3a/C3aR1 axis in regulating adipose thermogenesis. Our findings establish a newly discovered role of the alternative complement pathway in adipose thermogenesis and highlight sex-specific considerations in potential therapeutic targets for metabolic diseases.

## Introduction

Obesity remains a serious global public health problem that greatly increases the risk of type 2 diabetes (T2D) and other associated diseases, including cardiovascular disease and many types of cancer[1, 2]. Bariatric surgery, currently the most effective durable option for treating severe obesity carries risk of surgical complications, death, and reoperation[3]. Glucagon-like peptide 1 receptor (GLP-1R) and gastric inhibitory polypeptide 1 receptor (GIP-R) dual agonists are the most promising medical treatments for obesity to date, largely dependent on suppressing food intake[4, 5]. Adaptive thermogenesis, the production of heat by the body in response to stimuli such as cold, is considered a promising therapeutic approach to counteract obesity[6]. Thermogenic adipocytes in the brown and white adipose tissue depots are specialized cells that are responsible for adaptive thermogenesis through uncoupling protein 1 (UCP1) enhancing the leakage of protons across the mitochondrial membrane and other futile cycling pathways[7-9]. Beige adipocytes embedded in white adipose tissues have multilocular lipid droplets and can promote thermogenesis like brown adipocytes[9, 10]. Beige adipocytes can arise from both preadipocyte progenitors and pre-existing mature unilocular white adipocytes[9, 11-15]. The formation of beige adipocytes is activated by cold exposure, β-adrenergic stimulation, and other stimuli[9]. The detection of brown adipose tissue (BAT) in human adults has further piqued interest in enhancing energy expenditure by increasing the thermogenic activity of brown and beige adipocytes[16-18]. However, therapies to stimulate the thermogenic capacity of adipose tissues based on cold exposure or β-adrenergic agonists remain clinically challenging. Therefore, there is considerable interest in finding novel mechanisms that promote browning and enhance thermogenesis in adipose.

The complement system consists of many distinct complement components that interact with one another. As an important part of the innate immune system, complement plays a key role in the defense against common pathogens[19]. A growing body of research suggests that certain components of the complement system also play an important role in the regulation of metabolic disorders such as diabetes and insulin resistance[20-22]. A critical complement component, complement factor D, also known as adipsin, is mainly secreted by adipocytes. Adipocytes synthesize the major components of the alternative complement pathway, but the role complement plays in adipose tissue homeostasis is unknown. Adipsin controls the alternative complement pathway by catalyzing the production of the C3 convertase, which then cleaves C3 to generate C3a and C3b[23]. We previously showed that C3a acts to increase beta cell insulin secretion, an effect that is dependent on C3a receptor 1(C3aR1)[21, 24]. Several studies show significant though opposing roles of adipsin and C3aR1 on systemic glucose homeostasis in diet-induced obesity[20, 21, 25]. These results suggest that there may be different cell-specific effects of adipsin/C3aR1 on obesity and diabetes. C3aR1 is predominantly expressed on immune cells and was not previously known to be expressed by adipocytes. Furthermore, it is unknown whether the adipsin/C3a/C3aR1 axis is involved in the regulation of systemic energy homeostasis and adipose thermogenesis.

In the present study, we show that *Adipsin* deficiency increased the thermogenic program in white adipose tissue (WAT) under ambient conditions and with cold exposure. We find that adipocytes express the anaphylatoxin receptor C3aR1. We generated adipocyte-specific *C3aR1* knockout (Ad-*C3aR1*^*-/-*^) mice to elucidate the physiological roles of C3aR1 in adipocytes. Our study demonstrated that Ad-*C3aR1*^*-/-*^ male mice exhibited enhanced thermogenic gene expression in subcutaneous adipose tissue. Male Ad-*C3aR1*^*-/-*^ mice also showed increased heat production in thermogenic adipose tissue. Further *in vitro* studies showed that deletion of *C3aR1* in primary subcutaneous adipocytes from male mice enhanced thermogenic gene expression and mitochondrial respiration. Notably, we found sexual dimorphism in *Adipsin* and *C3aR1* gene expression. *Adipsin* was lower in both adipose tissue and thermogenic adipocytes of female mice compared to males. Subcutaneous adipose tissue *C3aR1* gene expression was also diminished in female mice than males. Furthermore, adipocyte-specific *C3aR1* knockout female mice are cold intolerant and their brown adipose tissues produce less heat compared to controls. Our results reveal the function of the adipsin/C3a/C3aR1 axis as a regulator of white adipose thermogenesis and energy metabolism in a sex-dependent manner.

## Results

### Adipsin Leads to Increased Energy Expenditure and Protection from Diet-induced Obesity

We previously found that adipsin positively regulates adipose tissue inflammation in mice fed a high fat diet (HFD) [21]. However, adipsin-deficient mice were mildly resistant to diet-induced obesity. To explore the potential role of adipsin in regulating whole-body energy homeostasis with metabolic stress, we compared the body weights of male wild type (WT) and *Adipsin*^*-/-*^ mice fed a regular diet (RD) or HFD. As expected, *Adipsin* mRNA levels were noticeably absent in all the fat tissues of the knockout (KO) mice (Figure S1A). While no change was seen in body weights between male WT and *Adipsin*^-/-^ mice (Figure S1B) on a regular diet, *Adipsin*^-/-^ mice displayed a mild-moderate but significant reduction in body weight on HFD compared to both WT and *Adipsin*^+/-^ mice (Figure 1A). Body composition analysis showed that the reduced weight gain of *Adipsin*^-/-^ mice was mainly due to decreased fat mass without significant differences between WT and *Adipsin*^+/-^ mice (Figure 1B). As body weight is determined by a balance between energy expenditure and energy intake, we assessed food intake and energy expenditure in *Adipsin*^-/-^ and control mice. Our results indicate that food intake was comparable between male WT and *Adipsin*^*-/-*^ mice (Figure 1C), suggesting that the reduced body weight gain in *Adipsin*^*-/-*^ mice was caused by increased energy expenditure. Indeed, *Adipsin*^*-/-*^ mice exhibited an increase in O_2_ consumption and CO_2_ production compared to WT mice (Figures 1D, 1E, S1C, and S1D).

**Figure 1.**
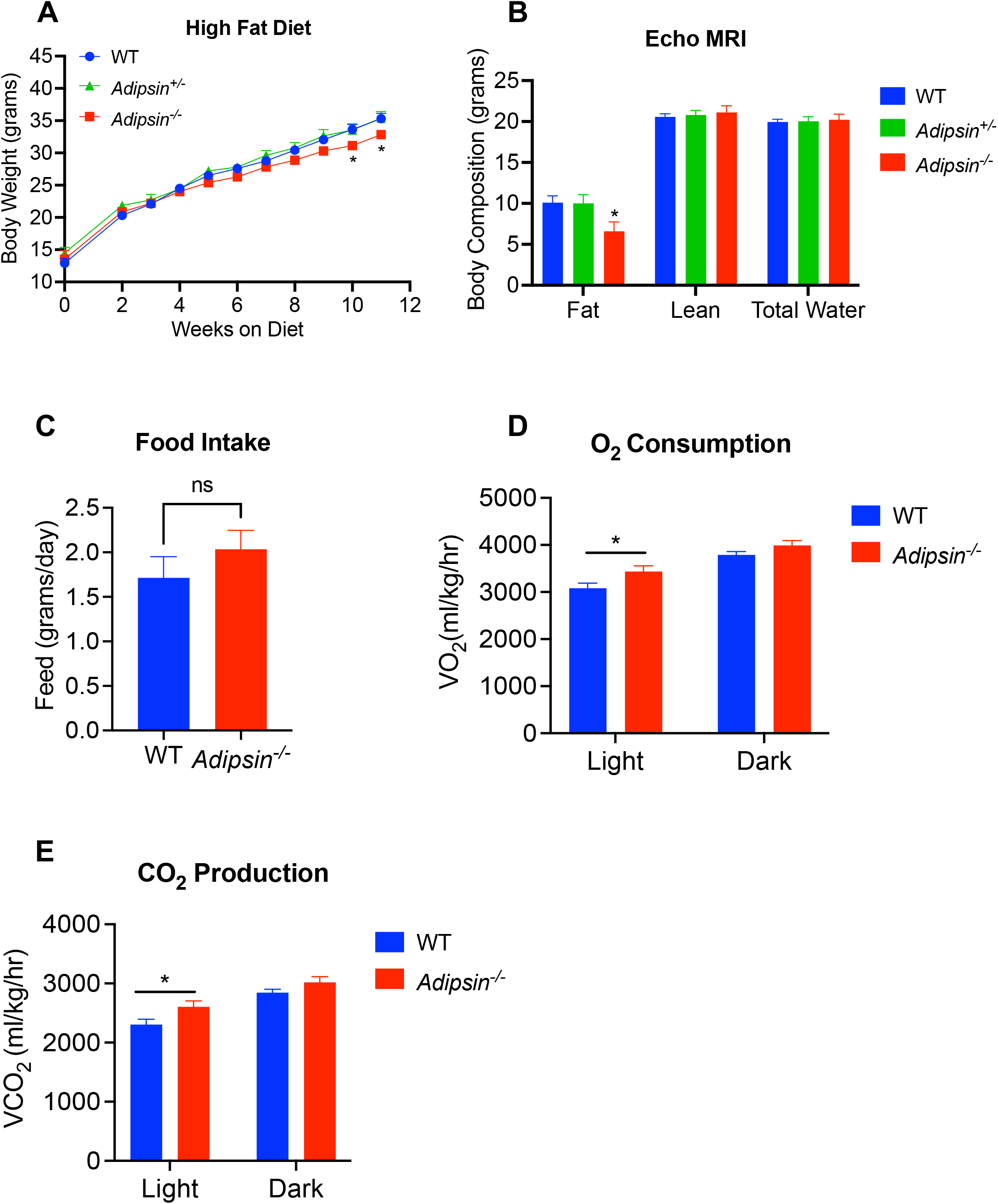
Mice deficient in Adipsin are protected from diet-induced obesity and display evidence of enhanced energy expenditure. **A)** Body weights of wild type (WT), *Adipsin* heterozygous and knockout male mice on high fat diet (HFD) for the indicated number of weeks. n = 8/group. **B)** Body composition of male mice from (A) after 12 weeks of HFD. n = 8/group. **C)** Food intake of male mice from (A) was measured daily after 4 weeks on HFD. n = 8/group. **D-E)** O_2_ consumption (**D**) and CO_2_ production (**E**) rates of WT and Adipsin knockout male mice were measured by indirect calorimetry using comprehensive lab animal monitoring system (CLAMS) after 4 weeks on HFD. n=8/group. Data are presented as mean± S.E.M. Unpaired two-tailed t test is used for comparison. n.s., not significant. *p < 0.05.

### Adipsin Deficiency Results in Enhanced Thermogenesis in White Fat

Since brown/beige adipocyte thermogenesis plays a key role in driving energy expenditure and adipsin is produced by adipocytes, we next examined whether there was enhanced adipose thermogenesis in *Adipsin*^*-/-*^ mice. Ablation of adipsin resulted in increased expression of key thermogenic genes *Ucp1, Ppargc1a, Ppargc1b*, and *Prdm16* at room temperature in visceral (VISC) white adipose tissue (WAT) (Figure 2A) with a similar trend of elevated *Ucp1* gene expression observed in subcutaneous (SubQ) WAT (Figure 2B). There was not a significant difference in thermogenic gene expression in the brown adipose tissue (BAT) between WT and *Adipsin*^*-/-*^ mice (Figure S2A).

**Figure 2.**
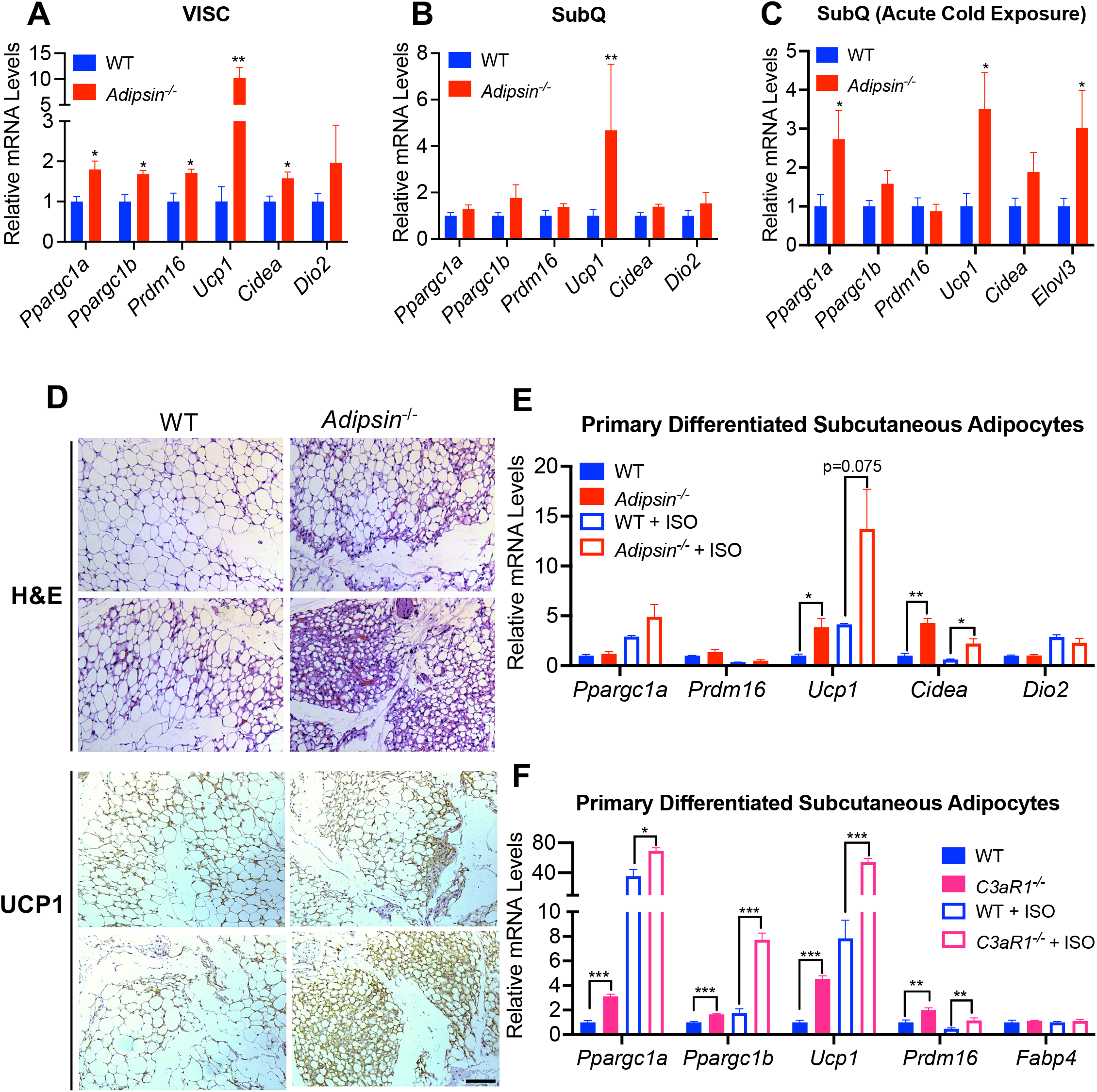
Adipsin deficiency promotes white adipose beiging. **A-B)** Thermogenic gene expression in visceral (VISC) (**A**) and subcutaneous (SubQ) (**B**) fat of 10-12 weeks old WT and *Adipsin* knockout male mice fed a regular diet at ambient temperature. n=5/group. **C)** Thermogenic gene expression in SubQ fat of 10-12 weeks old WT and *Adipsin* knockout male mice following an acute (6 hours) cold exposure. n=10/group. **D)** Hematoxylin and eosin and UCP1 immunohistochemistry staining of inguinal white adipose tissue sections from 10-weeek-old WT and *Adipsin* knockout male mice at ambient temperature and following acute cold exposure. Images are shown at 20x magnification. Scale bar, 200 μm. **E)** Thermogenic gene expression in primary subcutaneous adipocytes treated with isoproterenol from WT and *Adipsin* knockout mice. n=3/group. **F)** Thermogenic gene expression in primary subcutaneous adipocytes treated with isoproterenol from WT and *C3aR1* knockout mice. n=6/group. Data are presented as mean± S.E.M. Unpaired two-tailed t test is used for comparison. *p < 0.05, **p < 0.01, ***p < 0.001.

SubQ WAT is prone to browning upon cold stimulation or β-adrenergic agonists[26]. To further study the role of adipsin in regulation of white fat browning, we examined the effects of cold exposure on *Adipsin*^*-/-*^ and WT mice. *Ucp1* was induced by 3-fold in the SubQ and VISC WAT of *Adipsin*^*-/-*^ compared to WT mice following acute cold exposure (Figures 2C and S2B). However, the expression of thermogenic genes was not altered in the BAT of acute cold-exposed *Adipsin*^*-/-*^ mice (Figure S2C). Because of growing evidence that beige fat also uses alternative thermogenic pathways to dissipate energy in the form of heat[9], we next examined changes in *Ucp1*-independent thermogenic genes upon cold stimulation of SubQ fat. Overall, the *Adipsin*^*-/-*^ group had a trend of higher *Ucp1*-independent thermogenic gene expression than the WT group (Figure S2D). Consistent with the gene expression data, SubQ WAT from *Adipsin*^*-/-*^ compared to that from WT mice showed more UCP1+ cells and reduced lipid stores (Figure 2D). After 1 week of chronic cold exposure, we did not detect differences in thermogenic gene expression between WT and *Adipsin*^*-/-*^ mice (Figures S2E, S2F, and S2G).

Since adipsin and its products can potentially act on many different cell types, we next examined whether the increased numbers of thermogenic adipocytes in SubQ adipose tissue of *Adipsin*^-/-^ mice was cell autonomous. Adipocytes differentiated in vitro from the SubQ adipose depot from *Adipsin*^-/-^ mice also displayed a 4-5 fold increase in the expression of *Ucp1* and cell death inducing DFFA like effector A (*Cidea*) compared to controls (Figure 2E), suggesting an adipocyte cell-autonomous role for adipsin in thermogenesis. Furthermore, this increased expression of thermogenic genes in *Adipsin*^-/-^ cells was also enhanced by the β-agonist isoproterenol (Figure 2E).

Adipsin catalyzes the formation of the C3 convertase, which can then result in the cleavage of C3 into C3a and C3b[21, 23]. C3a acts on cells through its G-protein coupled receptor (GPCR) C3aR1 whereas C3b activates the C5 convertase that can result in formation of the C5b-C9 membrane attack complex [21, 27-29]. To determine whether adipsin regulates subcutaneous adipocyte browning acutely through its downstream C3a/C3aR1 signaling pathway, we next assessed primary subcutaneous adipocytes from WT and *C3aR1* whole-body KO mice. We found that subcutaneous adipocytes from *C3aR1* KO mice differentiated *in vitro* showed a robust increase in the expression of thermogenic genes compared to WT controls (Figure 2F). Stimulating these cells with isoproterenol further augmented thermogenic gene expression in both groups with *C3aR1* KO adipocytes remaining 3-5 fold higher than controls (Figure 2F). These data suggest that adipsin through C3a/C3aR1 signaling regulates a browning phenotype in subcutaneous adipocytes. Furthermore, we observed that *C3aR1*^*-/-*^ mice exhibited a trend towards mildly increased energy expenditure as measured by O_2_ consumption and CO_2_ production compared to WT mice (Figures S2H and S2I). Collectively, these results suggest that adipocyte C3aR1 may regulate adipose thermogenesis and systemic energy homeostasis.

### Adipocyte C3aR1 Regulates Subcutaneous Adipocyte Thermogenic Gene Expression and Mitochondrial Respiration

To directly interrogate the role of C3aR1 on adipocytes in adaptive thermogenesis, we generated adipocyte specific C3aR1 KO mice by crossing *C3ar1* floxed mice with Adiponectin-Cre transgenic mice (Ad-*C3aR1*^-/-^). Because of the cellular heterogeneity in adipose tissues with macrophages and immune cells expressing high levels of C3aR1, we analyzed *C3ar1* gene expression in the adipocyte fraction to confirm the adipocyte specific knockout in this system. Our results showed 50-65% knockdown of *C3ar1* in adipocytes in Ad-*C3aR1*^-/-^ compared to controls (Figure 3A). Male Ad-*C3aR1*^-/-^ mice displayed increased expression of thermogenic genes such as *Ppargc1a, Ppargc1b, Prdm16, Ucp1* and Cidea compared to control mice, mostly in SubQ adipose tissue (Figures 3B and S3B). Thermogenic gene expression in VISC fat of control and Ad-*C3aR1*^-/-^ mice showed no difference (Figure S3A). Furthermore, we observed a trend towards increased heat production in brown and SubQ adipose tissue of Ad-*C3aR1*^-/-^ mice compared to controls (Figures 3C and 3D).

**Figure 3.**
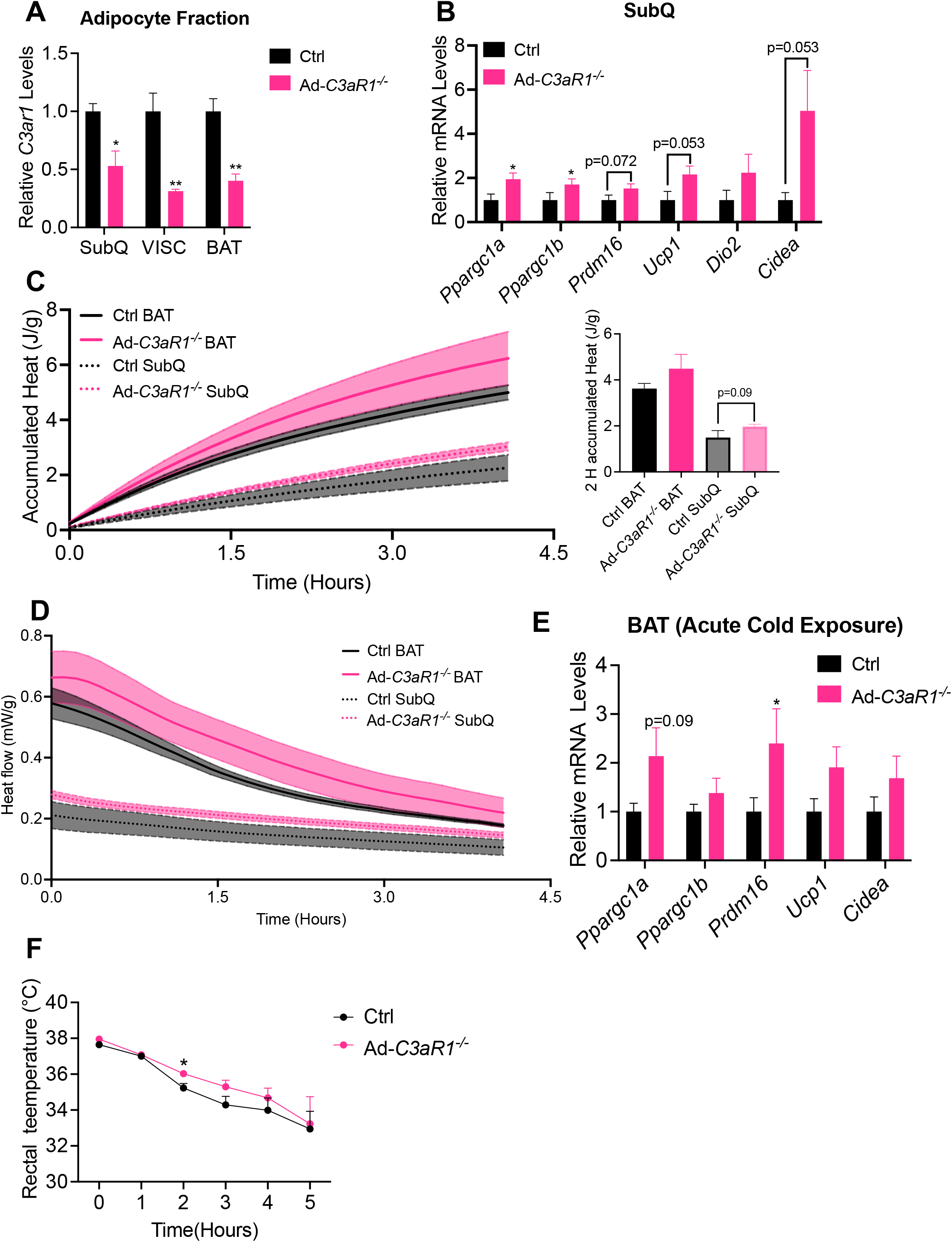
Adipocyte-specific C3aR1 knockout male mice and adipose thermogenic capacity. **A)** Relative *C3aR1* gene expression in the adipocyte fraction of 7 weeks old control and Ad-*C3aR1*^*-/-*^ male mice fed a regular diet at ambient temperature. n=3/group. **B)** Thermogenic gene expression in subcutaneous (SubQ) fat of 10-12 weeks old control and Ad-*C3aR1*^*-/-*^ male mice fed a regular diet at ambient temperature. n=4/group. **C-D)** Accumulated heat (**C**) in J and heat flow (**D**) in mW recorded from wells in duplicates containing adipose tissue from control and Ad-*C3aR1*^*-/-*^ male mice fed a regular diet at ambient temperature. n=4/group. **E)** Thermogenic gene expression in brown fat of 10-12 weeks old control and Ad-*C3aR1*^*-/-*^ male mice following an acute (6 hour) cold exposure. n=6/group. **F)** Rectal temperature of 10-12 weeks old control and Ad-*C3aR1*^*-/-*^ male mice during acute cold exposure. n=6/group. Data are presented as mean± S.E.M. Unpaired two-tailed t test is used for comparison. *p < 0.05, **p < 0.01.

In addition, when exposed to cold for 6 hours, we found a trend of higher expression of thermogenic genes in the SubQ and BAT of the adipocyte C3aR1 KO group compared to controls (Figures 3E and S3B). Male Ad-*C3aR1*^-/-^ mice also showed mild improvements in cold tolerance when compared to controls (Figure 3F). There was no difference in body weight or adipose tissue weight between the two groups (Figures S3C and S3D). In addition, thermogenic gene expression did not differ between chronic cold-exposed control and Ad-*C3aR1*^*-/-*^ mice (Figures S3E and S3F).

To exclude any potential developmental effects contributing to the increased thermogenic gene expression observed in SubQ WAT of Ad-*C3aR1*^-/-^ mice, we assessed beige adipocyte development and gene expression in vitro. We isolated cells from the stromal vascular fraction (SVF) of adipose tissues from control and Ad-*C3aR1*^-/-^ mice and then differentiated them to mature adipocytes. We found that Ad-*C3aR1*^-/-^ SubQ adipocytes showed a robust 4-fold increase in *Ucp1* and 2-fold increase in *Cidea* compared to controls (Figure 4A). Stimulation of these cells with isoproterenol further augmented thermogenic gene expression, especially in adipocytes lacking C3aR1 (Figure 4A). However, knockout of *C3aR1* in brown adipocytes did not significantly impact thermogenic gene expression (Figure 4B). These results suggest that C3aR1 on adipocytes regulates thermogenic gene expression mostly in SubQ but not brown adipocytes.

**Figure 4.**
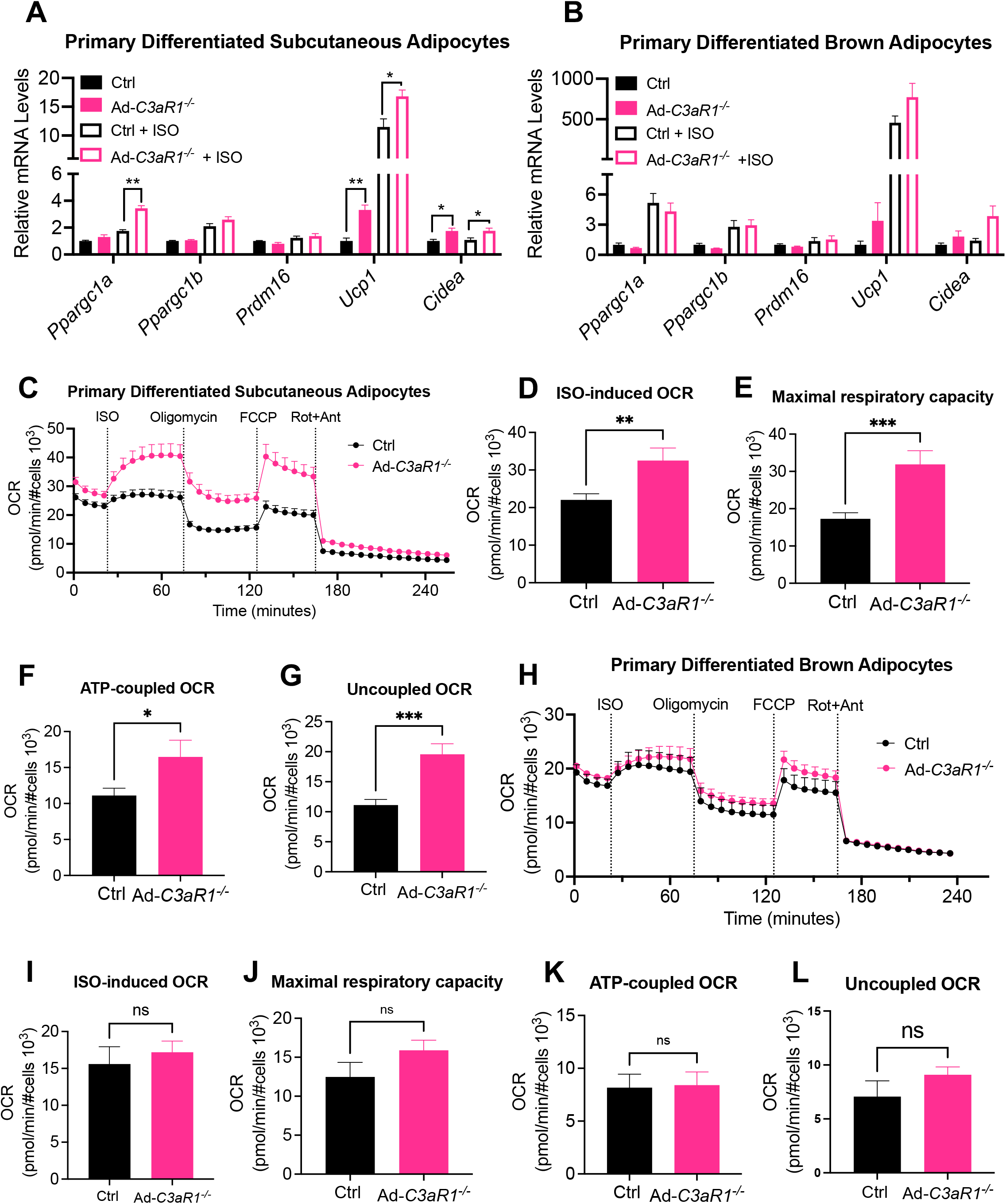
C3aR1 deficiency in male subcutaneous adipocytes results in increased thermogenesis and respiration *in vitro*. **A)** Thermogenic gene expression in primary subcutaneous adipocytes treated with isoproterenol from control and Ad-*C3aR1*^*-/-*^ male mice. n=4/group. **B)** Thermogenic gene expression in primary brown adipocytes treated with isoproterenol from control and Ad-*C3aR1*^*-/-*^ male mice. n=4/group. **C)** Oxygen consumption rate (OCR) of primary control and Ad-*C3aR1* knockout subcutaneous adipocytes. ISO: isoproterenol; FCCP: carbonyl cyanide 4 (trifluoromethoxy) phenylhydrazone; Rot: rotenone; Ant: antimycin. n=20/group. **D-G)** Quantification of isoproterenol-stimulated OCR (**D**), maximal respiratory capacity(**E**), Adenosine triphosphate (ATP)-coupled OCR (**F**), and uncoupled OCR (**G**) from primary control and Ad-*C3aR1* knockout subcutaneous adipocytes. **H)** OCR of primary control and Ad-*C3aR1* knockout brown adipocytes. n=20/group. **I-L)** Quantification of isoproterenol-stimulated OCR (**I**), maximal respiratory capacity(**J**), Adenosine triphosphate (ATP)-coupled OCR (**K**), and uncoupled OCR (**L**) from primary control and Ad-*C3aR1* knockout brown adipocytes. Data are presented as mean± S.E.M. Unpaired two-tailed t test is used for comparison. n.s., not significant. *p < 0.05, **p < 0.01, ***p < 0.001.

Compared with white adipocytes, mitochondria in beige adipocytes and brown adipocytes are significantly more numerous, larger, and contain more cristae[9]. The increased expression of thermogenic genes in *C3aR1* knockout SubQ adipocytes prompted us to investigate the respiratory activity of isolated adipocytes from control and adipocyte specific *C3aR1* KO mice. Subcutaneous adipocytes from Ad-*C3aR1*^-/-^ mice displayed double the maximal respiratory capacity and 50% more isoproterenol-induced respiration that was mostly due to uncoupled respiration but also with elevated Adenosine triphosphate (ATP)-coupled respiration compared to controls (Figures 4C, 4D, 4E, 4F and 4G). However, Seahorse analyses showed no significant differences in brown adipocyte respiration between the control and the *C3aR1* knockout group (Figures 4H, 4I, 4J, 4K, and 4L). These data suggest that loss of *C3aR1* on beige adipocytes stimulates both uncoupled and ATP-coupled respiration as a mechanism to enhance thermogenic capacity in subcutaneous adipocytes.

### Sex-associated Differences in the Alternative Complement Pathway

Clinical studies have illuminated sex differences in alternative complement pathway activity but not in the classical and lectin pathways[30]. The circulating levels of two important alternative complement components, C3 and adipsin, show sex differences[30, 31]. These sex differences in humans prompted us to investigate if there is a similar sexual dimorphism in mice. We found no difference in circulating adipsin levels between male and female mice (Figure 5A). Since adipsin is predominantly secreted by adipose tissues, we assayed *Adipsin* and *C3aR1* levels in adipose tissues of male and female mice. We found that *Adipsin* mRNA level is lower in the thermogenic adipose tissue of female mice, especially in SubQ adipose tissue (Figures 5B, 5C, and 5D). However, *C3aR1* was only decreased in the SubQ adipose tissue of female mice compared with males (Figure 5B). *C3aR1* gene expression was not different between female and male mice in the visceral adipose tissue and BAT (Figures 5C and 5D). Next, we assessed *Adipsin* mRNA levels in the adipocyte fractions from the different major fat depots. Our results show that *Adipsin* was significantly lower in subcutaneous and brown adipocytes of female mice compared with males (Figure 5E). These results reveal that there is sexual dimorphism in *Adipsin* expression in SubQ and brown but not visceral adipocytes in mice.

**Figure 5.**
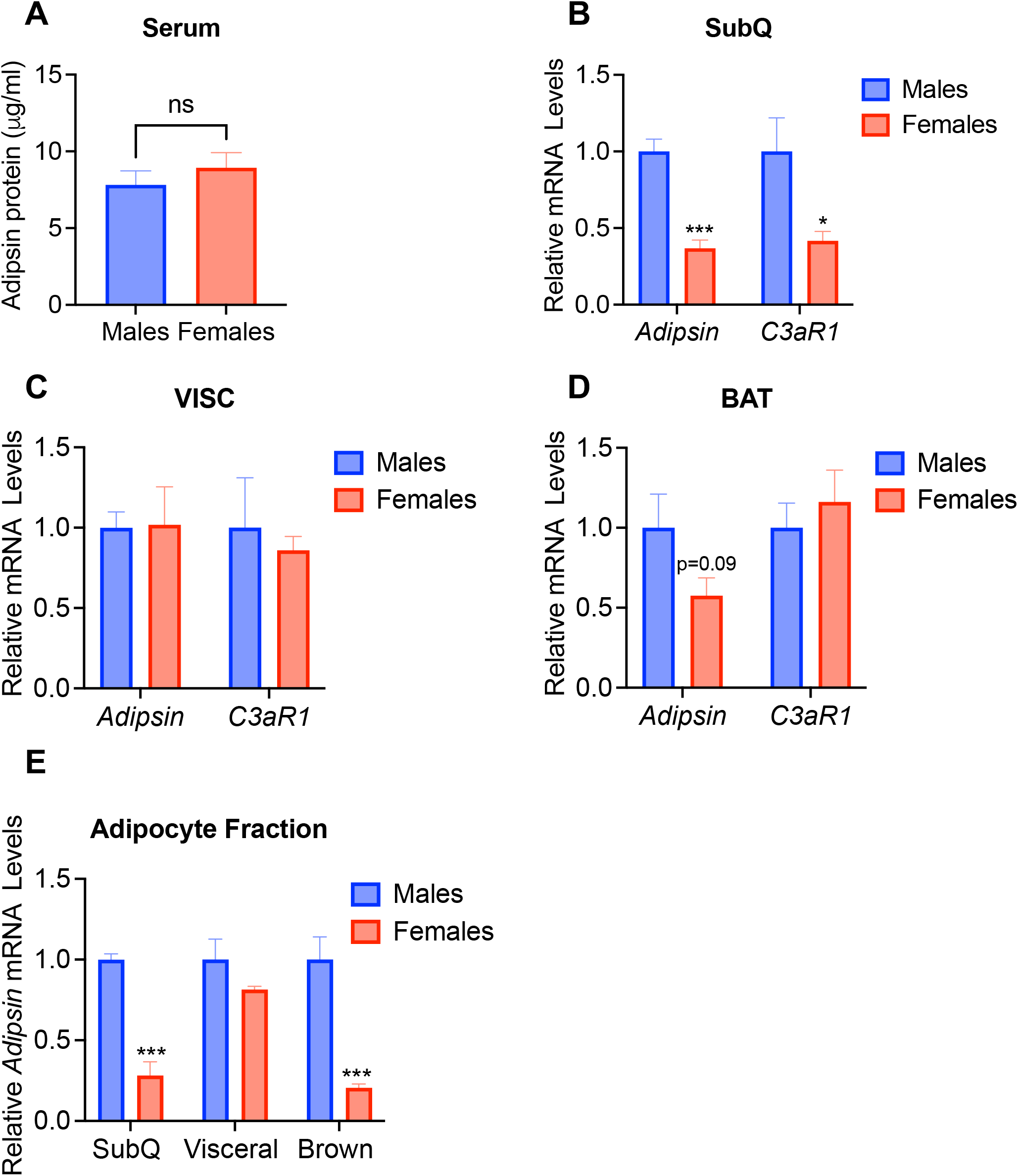
Sex-dependent differences in alternative complement pathway components. **A)** ELISA for adipsin from serum of wild type (WT) mice at 10-11 weeks old on regular diet. n=8/group. **B-D)** Relative *Adipsin* and *C3aR1* gene expression in subcutaneous (SubQ) (**B**), visceral (VISC) (**C**) and brown (**D**) adipose tissue of 10-11 weeks old WT mice on regular diet. n=5/group. **E)** Relative *Adipsin* gene expression in the adipocyte fraction of 10-11 weeks old WT mice on regular diet. n=5/group. Data are presented as mean± S.E.M. Unpaired two-tailed t test is used for comparison. n.s., not significant. *p < 0.05, ***p < 0.001.

### Ad-*C3aR1*^*-/-*^ Female Mice Display Decreased Thermogenic Gene Expression and Cold Intolerance

Since we observed differences in expression of alternative pathway components between male and female mice, we tested if adipocyte C3aR1 expression may have different effects in female mice. In stark contrast to male mice, Ad-*C3aR1*^-/-^ female mice exhibited substantial impairments at defending their core body temperature during acute cold stress compared to controls (Figure 6A). We found *Ppargc1b* to be decreased and similar trends for *Prdm16* and *Ucp1* in the BAT of Ad-*C3aR1*^*-/-*^ female mice compared to the control group during acute cold stress (Figure 6B). *Ucp1* also trended towards being lower in Ad-*C3aR1*^*-/-*^ female knockout mice compared to controls at ambient temperature (Figure 6C). There was a similar trend of decreased *Ucp1, Ppargc1a*, and *Cidea* in the brown fat of Ad-*C3aR1*^-/-^ knockout mice compared to controls (Figures S4A and S4B). Importantly, we observed 30% less heat production in the BAT of Ad-*C3aR1*^-/-^ female mice compared to controls (Figures 6D and 6E). These results suggest that impaired BAT thermogenesis in Ad-*C3aR1*^-/-^ female mice significantly compromised their endothermic response to acute cold exposure. There was no difference in the body weights and thermogenic adipose tissue depots between the two groups after acute cold exposure (Figures S4C and S4D). We did not detect any differences in brown adipocyte morphology and lipid stores in the BAT between the two groups (Figure S4E). Consistent with the changes in thermogenic gene expression, UCP1 staining was also fainter in the BAT of female Ad-*C3aR1*^-/-^ mice compared to controls (Figure S4F). Collectively, these results suggest that adipocyte C3aR1 regulates thermogenic adipose tissue function in a sex-dependent manner.

**Figure 6.**
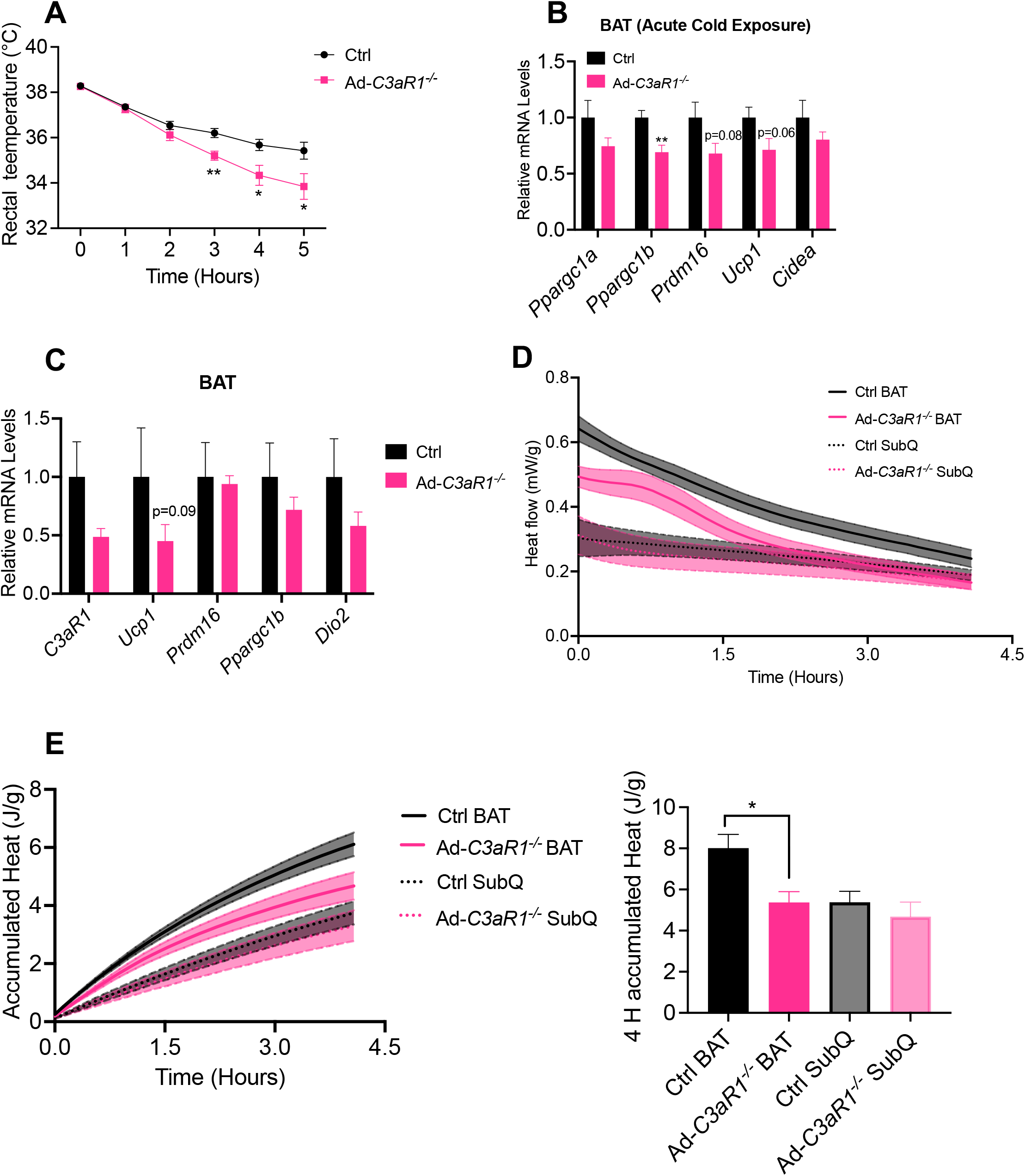
Adipocyte-specific *C3aR1* knockout female mice have impaired thermogenic capacity and are cold intolerant. **A)** Rectal temperature of control and Ad-*C3aR1* knockout female mice during acute cold exposure. n=12/group. **B)** Thermogenic gene expression in brown adipose tissue (BAT) of 10-12 weeks old control and Ad-*C3aR1*^*-/-*^ female mice following an acute (6 hour) cold exposure. n=6/group. **C)** Thermogenic gene expression in BAT of 10-12 weeks old control and Ad-*C3aR1* knockout female mice fed a regular diet at ambient temperature. n=6/group. **D-E)** Heat flow (**D**) in mW and accumulated heat (**E**) in J and recorded from wells in duplicates containing adipose tissue from control and Ad-*C3aR1*^*-/-*^ female mice fed a regular diet at ambient temperature. n=4/group. Data are presented as mean± S.E.M. Unpaired two-tailed t test is used for comparison. *p < 0.05, **p < 0.01.

## Discussion

Adipose tissues can synthesize the major components of the alternative complement pathway and are also one of the targets of complement activation in an autocrine or paracrine fashion[23]. Although alternative complement activation is thought to promote the pathological progression of obesity-related metabolic diseases, the precise physiological mechanisms linking alternative complement pathways to metabolic dysfunction are not well defined. Here, we provide an essential role of the alternative complement adipsin/C3a/C3aR1 signaling axis in the cold-induced browning of white adipocytes. Deletion of *Adipsin* or *C3ar1* increases subcutaneous adipocyte thermogenic gene expression, resulting in adipocyte browning in cold-exposed male mice. Importantly, we also demonstrate sexual dimorphism in the expression of *Adipsin* and *C3ar1*, key alternative complement components in adipose tissues. Critically, the effect of adipocyte specific C3aR1 deficiency on adipocyte browning are diametrically opposite between male and female mice. Here we provide a rare molecular example of a non-sex hormone that regulates thermogenic adipose function in a sex-dependent fashion. Although it is unclear how much sex differences have been ignored in the past, our study highlights the importance of assessing for potential sexual dimorphism in adipose biology. C3a/C3aR1 seems to inhibit browning in males while stimulating adipocyte thermogenesis in female mice. This could be due to different tonicity of C3aR1 signaling between sexes, other external inputs, and/or the intrinsic differences in the C3aR1 signaling pathway between male and female beige adipocytes. How the downstream C3aR1 signaling pathway in adipocytes may differ between male and female mice will be of great future interest. It is noteworthy that autoimmune complement diseases in humans also display sexual dimorphism[31].

External stimuli can be leveraged to offset metabolic diseases by activating thermogenesis in brown and beige adipose tissues[32]. Adipsin is the rate-limiting enzyme in the alternative pathway and controls downstream products such as C3a, C3b, and C5a[33]. Our data suggest that the SubQ beige adipocytes are a major target of adipsin/C3a through C3aR1. During complement activation, adipsin catalyzes the formation of the C3 convertase. C3 is highly abundant, but C3a is quickly converted by carboxypeptidase B or N into the inactive C3a-desArg45 that does not signal through C3aR1[20, 34]. Indeed, the data here suggest that *C3aR1* deficiency significantly upregulated thermogenesis and mitochondrial respiration in male subcutaneous adipocytes. On the other hand, C3a-C3aR1 signaling is also involved in macrophage-mediated inflammation[25]. Alternatively activated macrophages and eosinophils have also been shown to enhance adipocyte browning and protect against diet-induced insulin resistance[35, 36]. Therefore, it would be of future interest to determine whether the C3a-C3aR1 signaling is involved in macrophage-related adipocyte browning in adipose tissues. Clinical studies have demonstrated sex differences in alternative complement activity and complement component levels in healthy Caucasians[30]. Alternative pathway activity is higher in males than in females[30]. Circulating levels of C3 are higher in males, while adipsin levels are higher in females[30]. Other clinical studies have also confirmed that C3 has higher expression in the cerebrospinal fluid and plasma of men than in women[31, 37]. Our results show for the first time that in C57BL/6J mice, circulating levels of adipsin do not differ by sex, but its gene expression is higher in white adipose tissue and thermogenic adipocytes in males than in females. These results suggest that biological sex should be taken into account in alternative complement-related pathology as well as in complement-targeted therapies. Both genetics (such as the X chromosome) and hormonal differences are known to explain the effects of sexual dimorphism on immunity. But what is causing the difference in the key complement components in the alternative pathway is still unclear. Several clues have suggested that sex may form a possible confounding factor in complement-mediated diseases. For example, in one age-related macular degeneration study, significant differences in alternative pathway component levels were found between the sexes[38]. Therefore, further research is needed to elucidate the importance of sexual dimorphism on the complement system. In the future, it will be critical to demonstrate how sex hormones affect the complement system by administrating estrogen and testosterone into healthy and castrated mice of both sexes. Alternative complement levels and activity can also be studied in human study participants receiving hormone replacement therapy.

Although complement factors are best known for treating autoimmune diseases, their effects on adipose thermogenesis are new to the adipose metabolism field. Our analysis reveals that adipocyte-specific knockout of *C3aR1* mice results in major sex differences in response to cold stimulation. In particular, female mice with *C3aR1* KO in adipocytes exhibit pronounced cold intolerance. We do not yet have a mechanistic understanding of the downstream signaling pathway of C3aR1 in regulating the thermogenic gene program in adipocytes and why the results are discrepant between males and females. This may include differences in mitochondrial mass or activity and/or sensitivity to acute and chronic thermogenic stimuli. To our knowledge, this is the first demonstration of sexual dimorphism for beige adipocytes where a molecular receptor shows diametrically opposed functions between sexes. Previous studies have also found sexual dimorphism in the effects of adipose thermogenesis or thermogenesis-related markers[34, 39]. Filatov et al.[39] found that the thermogenic protein UCP1 was induced by cold exposure at the mRNA and protein levels in male Pacap-null mice; however, UCP1 protein induction was attenuated after cold adaptation in female Pacap-null mice. The observed sex-specific differences in the effects of C3aR1 on fat thermogenesis may be related to estrogen. There is evidence for sexual dimorphism in regulation of brown adipose tissue function and adipose tissue browning in rodent models and humans[40, 41]. For example, Benz et al. demonstrated that estrogen receptor α signaling in adipocytes can directly activate lipolysis[42]. In future studies, it will be of interest to determine if adipocyte C3aR1 is involved in estrogen receptor α signaling-related lipolysis in adipocytes. The mechanistic relationship between estrogen and complement factors remains to be understood with potential for crosstalk between the pathways. Our study highlights that the physiological response to cold stress is regulated differently between the sexes and stresses the importance of analyzing both sexes when assessing mechanisms of energy metabolism and fat thermogenesis.

## Materials and Methods

### Mice

WT and *Adipsin*^*-/-*^ mice were backcrossed to C57BL/6J background as described previously[21]. Whole body C3aR1 KO mice were purchased from Jackson laboratories (Strain #005712). *C3ar1* floxed mice are on the C57Bl/6J background as described[43]. Adipoq-Cre mice were purchased from Jackson laboratories (Strain #:028020). *C3ar1* floxed homozygous mice were used in the experiments as controls from the same backcross generation. All mice were maintained in plastic cages under a 12/12-h light/dark cycle at constant temperature (22°C) with free access to water and food. All animal studies were approved by the Institutional Animal Care and Use Committee and Research Animal Resource Center at Weill Cornell Medical College. For the diet-induced obesity model, mice were fed a 60% HFD (D12492i, Research Diets) for 16 weeks.

### Cold-induced Thermogenesis

All the experimental mice were placed in a cold chamber (4°C) for 6 hours (acute cold exposure) or 1 week (chronic cold exposure) with free access to water and food. During acute cold exposure, rectal temperature was measured using a BAT-12 Microprobe Thermometer (Physitemp, NJ, USA).

### Indirect Calorimetry

Metabolic rate was measured by indirect calorimetry in metabolic cage, a component of the Comprehensive Lab Animal Monitoring System (CLAMS; Columbus Instruments). Mice were housed individually and maintained at 22°C under a 12/12-h light/dark cycle. Food and water were available ad libitum.

### Histological Analysis

The adipose tissue was immediately perfused with PBS and fixed with 10% neutral-buffered formalin (VWR). Adipose tissues were then transferred to 70% ethanol. Paraffin embedding, sectioning and the hematoxylin and eosin (H&E) staining were done by the MSKCC Zuckerman Research Center Laboratory of Comparative Pathology core facility. For UCP1 immunohistochemistry, slides were dewaxed in xylene, hydrated in 100%, 95%, 80% and 70% ethanol, and rinsed in water, and antigen retrieval was performed in 10 mM sodium citrate buffer (pH=6.0) by boiling sections. Quenching of endogenous peroxidases was performed using 3% H_2_O_2_ solution (VWR). Slides were incubated with rabbit polyclonal UCP1 antibody (Abcam, ab10983) overnight at 4°C. Slides were washed in PBST and followed by incubation with HRP-conjugated secondary antibody (goat antirabbit, VWR/Jackson Immunoresearch). Then avidin–biotin peroxidase complex (Vectastain ABC kit, PK-7200, Vector Laboratories) was added for 30 min. Slides were developed using 3,3’-diaminobenzidine or 3-amino-9-ethyl carbazole (Sigma) and counterstained with hematoxylin.

### *In vitro* differentiation of primary adipocytes

For primary adipocytes, SVF from inguinal or visceral adipose from 6-to 7-week-old mice was prepared and differentiated for 6–8 days. Primary white adipocytes were cultured in DMEM/F12K media (Gibco) with 10% fetal bovine serum (FBS) at 37C°C, 5% CO2, until confluent. White adipocytes were differentiated via a 48 hours treatment with 0.5 mM 3-isobutyl-1-methylxanthine (IBMX), 1 mM dexamethasone, 850 nM insulin and 1 mM rosiglitazone, followed by 48 hours with 850 nM insulin and 1 mM rosiglitazone, and a further 48 hours with 850 nM insulin. Primary brown adipocytes were cultured in DMEM/F12K (Gibco) with 10% FBS at 37C°C, 5% CO_2_, until confluent. Brown adipocytes were differentiated via a 48 hours treatment with 0.5 mM IBMX, 5 μM dexamethasone,1 nM T3, 2 μM tamoxifen, 400 nM insulin and 1 μM rosiglitazone, followed by 48 hours with 400 nM insulin and 1 μM rosiglitazone, and a further 48 hours with 400 nM insulin. The fully differentiated adipocytes were treated with isoproterenol (100 nM) or C3a (100 nM from R&D) for 6 hours.

### Seahorse Assay

Primary subcutaneous adipocytes and brown adipocytes were plated into 96 well cell culture microplates (Agilent) and cells were plated in seahorse XF base medium (Agilent) containing 2mM L-glutamine and 5mM glucose adjusted to pH7.4. The seahorse microplate was incubated at in an incubator without supplemental CO_2_ at 37°C for 1 hour before assay. Oxygen consumption rates and extracellular acidification rates were measured using a XFe96 Seahorse (Agilent Technologies, Santa Clara, CA, USA). 0.5 μM rotenone and antimycin, 1 μM isoproterenol, 3 μM oligomycin and 2 μM carbonyl cyanide 4-(trifluoromethoxy)phenylhydrazone (FCCP) were injected during the assay.

### RNA Extraction and Real-Time Quantitative PCR

Total RNA from adipose and adipocytes were isolated using the RNeasy Mini kit (Qiagen) per manufacturer’s protocol. 1 μg RNA was reverse transcribed using high-capacity cDNA RT kit (Thermo). qPCR was performed using the SYBR Green Master Mix (Quanta) and specific gene primers on QuantStudio 6 Flex system (Thermo Fisher Scientific). Relative mRNA levels were determined by normalized to Ribosomal Protein S18 (Rps18) levels using the Delta-delta-cycle threshold (ΔΔCT) method. Primer sequences are listed in Supplementary Table 1.

### Blood adipsin measurements

For measuring mouse serum adipsin levels, blood was collected from the tail vein, the serum was separated by centrifugation and then adipsin levels were measured by ELISA (R&D) following the manufacturer’s protocol.

### Isothermal microcalorimeter assay

For heat detection experiments, we used the Cal-ScreenerTM a 48-channel isothermal microcalorimeter (Symcel SverigeAB, Spånga, Sweden), with its corresponding 48-well plate (calPlateTM). Each well consists of a screw-capped titanium vial. Data was continuously collected with the corresponding calViewTM software (Version 1.0.33.0, 2016, Symcel Sverige AB). For the assays, the machine was set and calibrated at 37 °C. General handling and device manipulation were done according to the manufacturer’s recommendations. We added 400 μL DMEM/F12K media (Gibco) with 10% fetal bovine serum and around 30-40 mg adipose tissue in each vial. The calView software was used for the calScreener IMC data collection and analysis. The direct measurement in IMC is heat flow (power in J/s) as a function of time. The heatflow gives the kinetic behavior and response of the sample over time. Data can also be expressed as the total accumulated heat (energy expressed in J) over time as an alternative data presentation of the cellular response to treatment.

### Statistics

All statistical analyses were performed using GraphPad Prism9. Unpaired two-tailed Student’s t tests and two-way ANOVA were used. Each in vitro experiment was repeated at least three times. p < 0.05 was considered statistically significant. The number of replicate samples presented in the text are biological replicates.

## Acknowledgments

We thank Dr. Peter S. Heeger from Cedars-Sinai for generously providing the *C3ar1* floxed mice. We thank Dr. Bruce M. Spiegelman for his advice on the project. L.M. was supported by a CSC Scholarship (201906050127). A.G. is supported by an ADA postdoctoral fellowship. This work was supported by the following grants: NIH R01 DK121140 (J.C.L.), R01 DK121844 (J.C.L.), and R01DK126944 (S.M.R.). The views expressed in this manuscript are those of the authors and do not necessarily represent the official views of the National Institute of Diabetes and Digestive and Kidney Diseases or the National Institutes of Health.

## Author Contributions

L.M. and J.C.L. designed the study and wrote the manuscript with input from all authors. L.M., A.G., A.R.N., E.C.A., S.M.R. and A.L. performed and analyzed the animal experiments. L.M., A.G., and A.R.N. developed and analyzed the *in vitro* experiments. S.M.R. and L.T. provided scientific input. J.C.L. conceived and supervised the study.

## Supplemental Figure Legends

**Figure S1.**
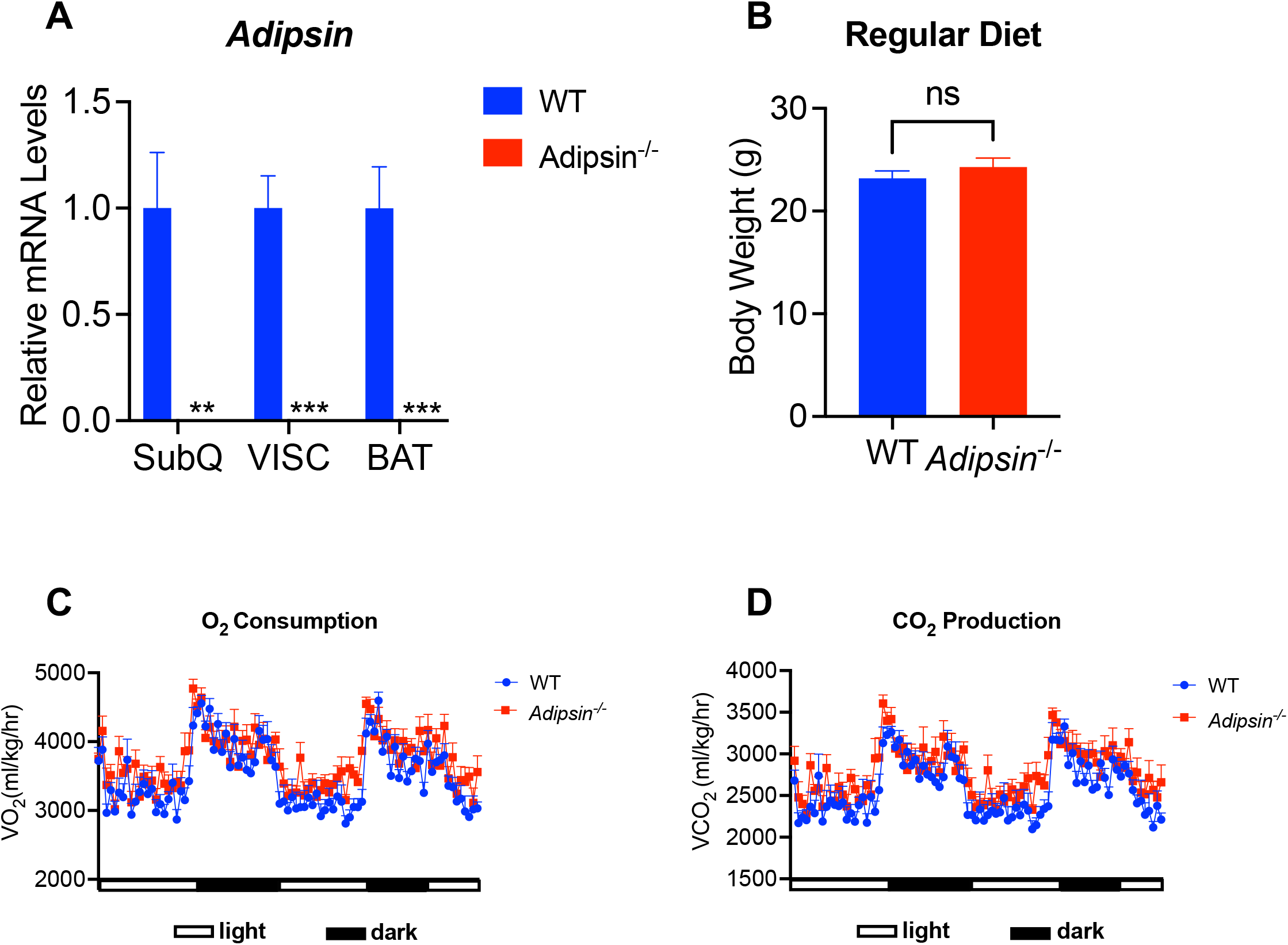
**A)** *Adipsin* mRNA expression in adipose tissues of 10-12 weeks old wild type (WT) and *Adipsin* knockout male mice fed a regular diet at ambient temperature. n=5**/**group. **B)** Body weights of WT and *Adipsin* knockout mice on 12 weeks of regular diet. n= 8/group. **C-D)** O_2_ consumption (**C**) and CO_2_ production (**D**) rates of WT and *Adipsin* knockout mice were measured by indirect calorimetry after 4 weeks on high fat diet (HFD). n=8/group. Data are presented as mean± S.E.M. Unpaired two-tailed t test is used for comparison. n.s., not significant. **p < 0.01, ***p < 0.001.

**Figure S2.**
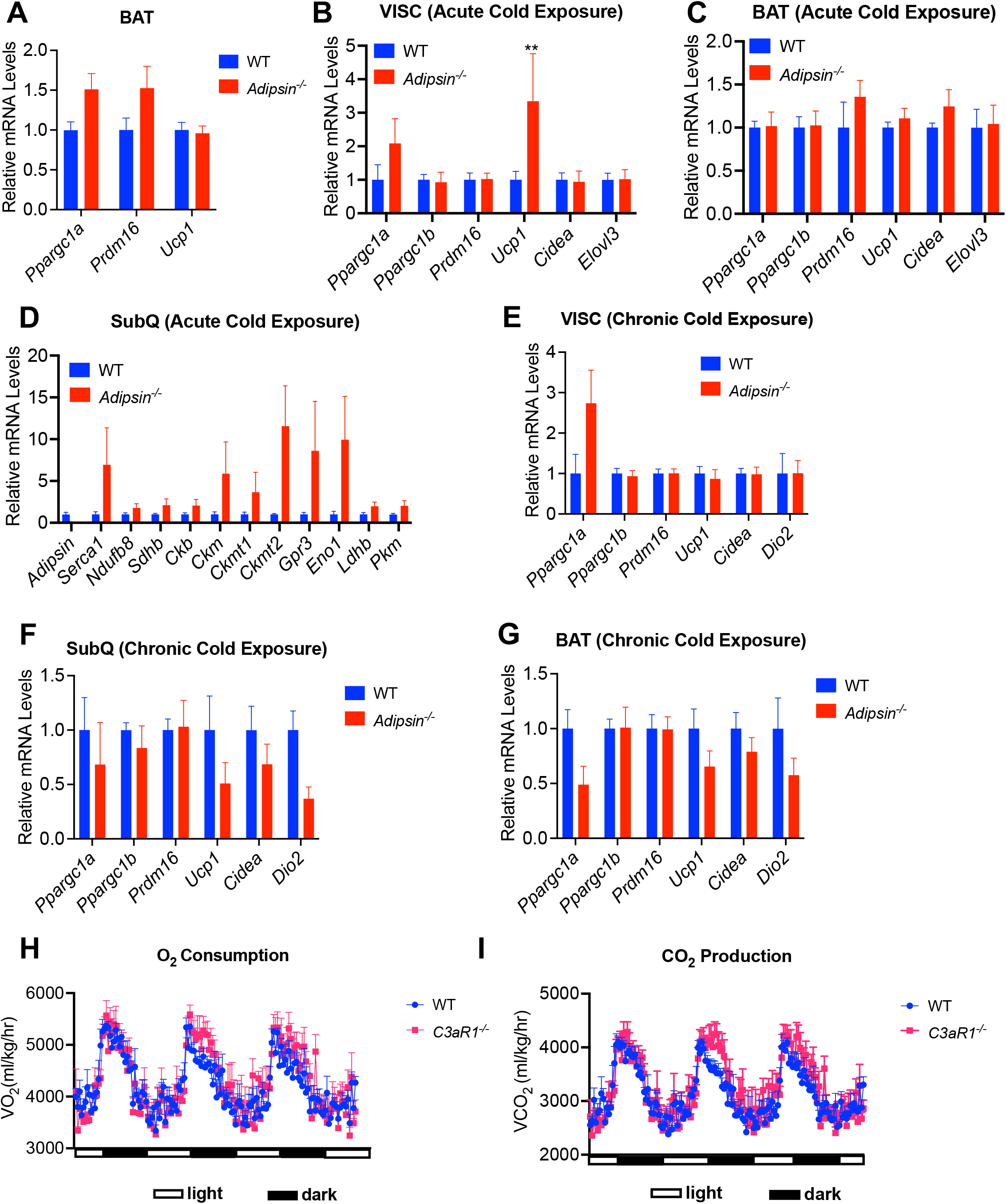
**A)** Thermogenic gene expression in brown adipose tissue (BAT) of 10-12 weeks old wild type (WT) and *Adipsin* knockout male mice fed a regular diet at ambient temperature. n=5/group. **B-C)** Thermogenic gene expression in visceral (VISC) (**B**) and BAT (**C**) of 10-12 weeks old WT and *Adipsin* knockout male mice following an acute (6 hour) cold exposure. n=5/group. **D)** UCP1-independent thermogenic gene expression in subcutaneous (SubQ) fat of 10-12 weeks old WT and *Adipsin* knockout male mice following an acute (6 hour) cold exposure. n=5/group. **E-G)** Thermogenic gene expression in VISC (**E**), SubQ (**F**) and brown (**G**) fat of 10-12 weeks old WT and *Adipsin* knockout male mice following a chronic (1 week) cold exposure. n=5/group. **H-I)** O_2_ consumption (**H**) and CO_2_ production (**I**) rates of WT and *C3ar1* knockout mice were measured by indirect calorimetry after 4 weeks on high fat diet (HFD). n=5/group. Data are presented as mean± S.E.M. Unpaired two-tailed t test is used for comparison. n.s., not significant. **p < 0.01.

**Figure S3.**
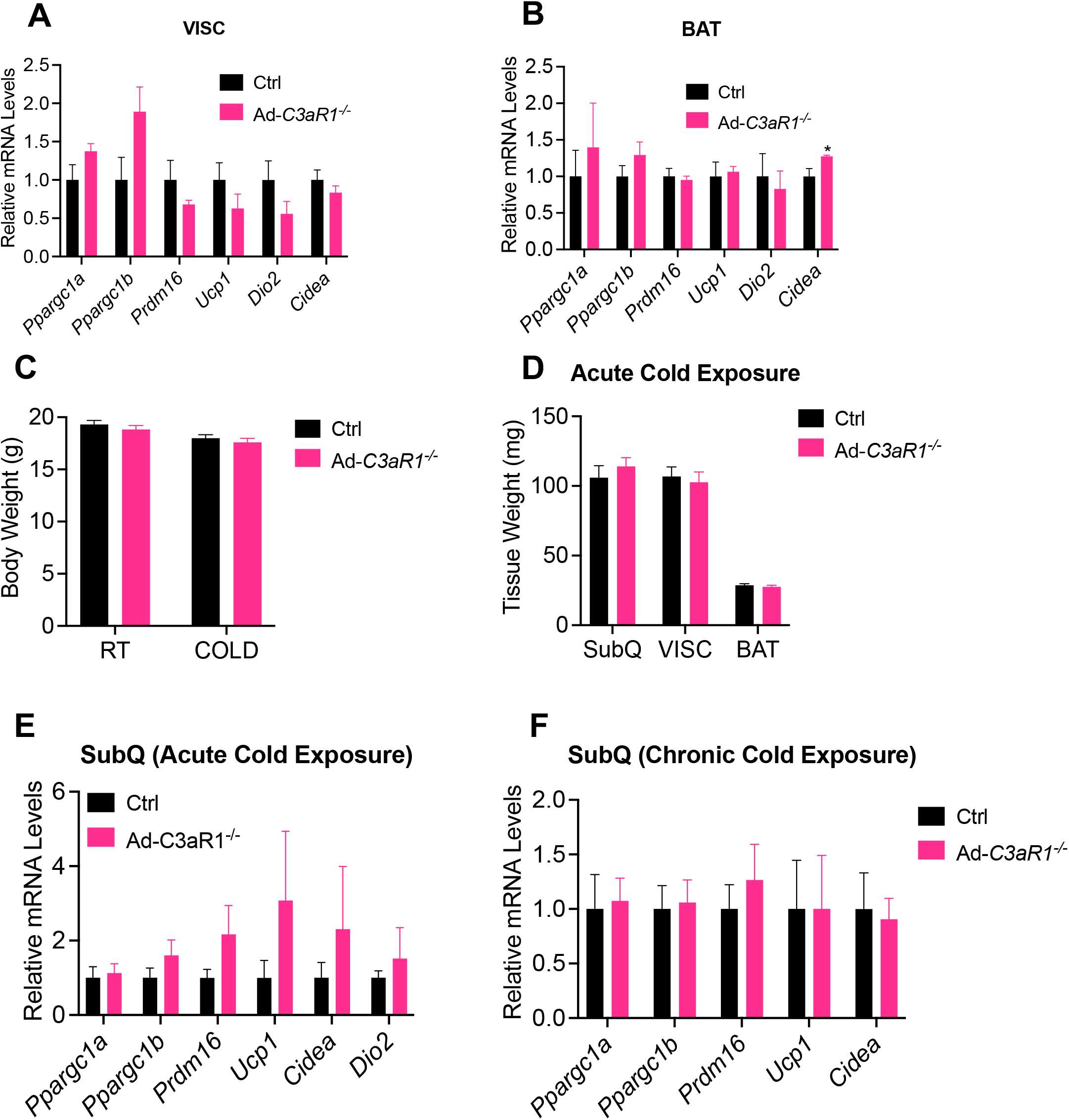
**A-B)** Thermogenic gene expression in visceral (VISC) (**A**) and brown (**B**) fat of 10-12 weeks old control and Ad-*C3aR1* knockout male mice fed a regular diet at ambient temperature. n=4/group. **C)** Body weights of control and Ad-*C3aR1* knockout male mice at room temperature and 6 hours post cold exposure. n=6/group. **D)** Adipose tissue weights of control and Ad-*C3aR1* knockout male mice 6 hours post cold exposure. n=6/group. **E)** Thermogenic gene expression of subcutaneous (SubQ) fat in control and Ad-*C3aR1* knockout male mice following an acute (6 hour) cold exposure. n=6/group. **F)** Thermogenic gene expression of SubQ fat in control and Ad-*C3aR1* knockout male mice following chronic (1 week) cold exposure. n=5/group. Data are presented as mean± S.E.M. Unpaired two-tailed t test is used for comparison.

**Figure S4.**
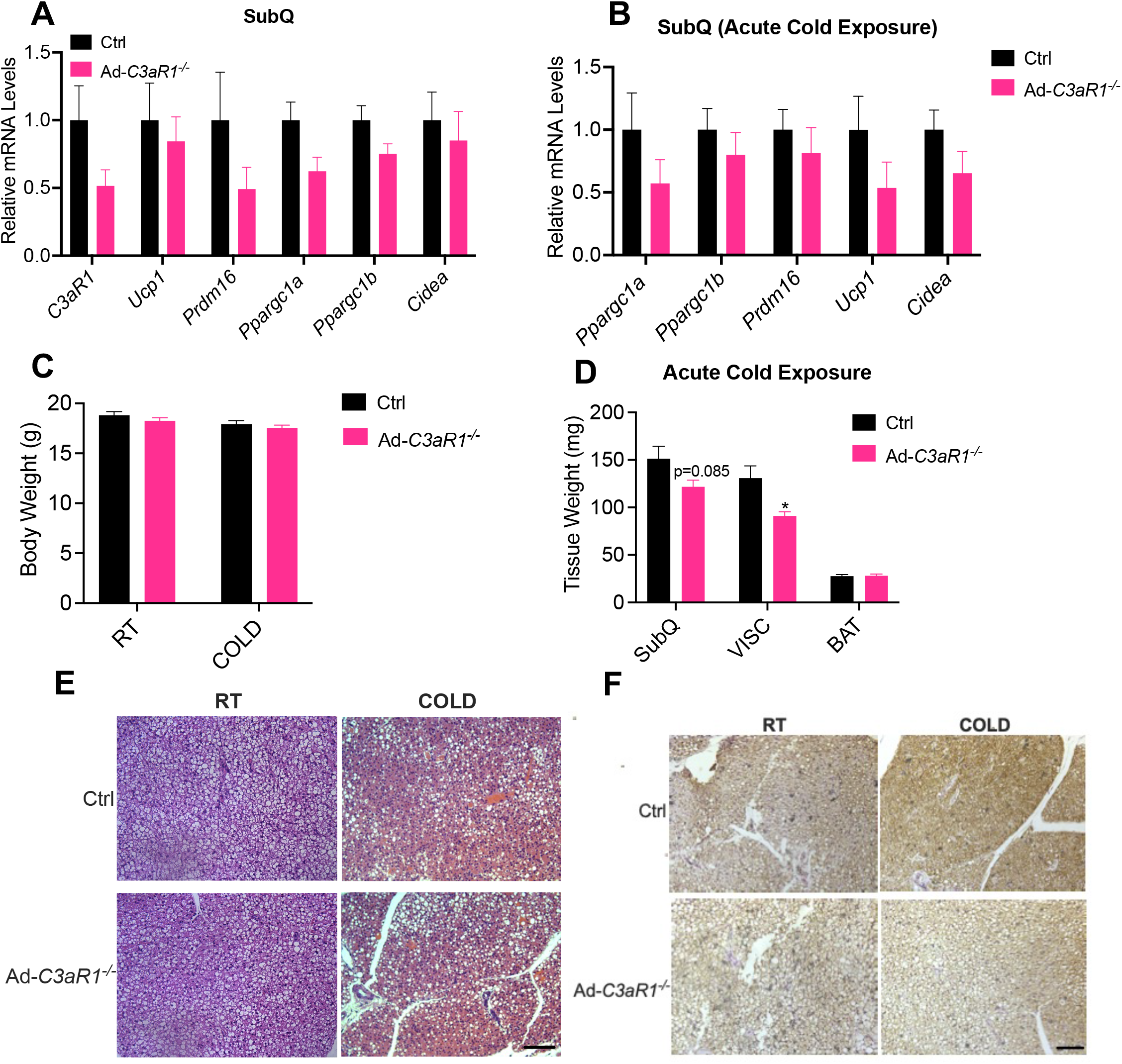
**A)** Thermogenic gene expression of subcutaneous (SubQ) fat in 10-12 weeks WT and Ad-*C3aR1* knockout female mice fed a regular diet at ambient temperature. n=6/group. **B)** Thermogenic gene expression of subcutaneous (SubQ) fat in 10-12 weeks WT and Ad-*C3aR1* knockout female mice following an acute (6 hour) cold exposure. n=6/group. **C)** Body weight of control and Ad-*C3aR1* knockout female mice at room temperature and 6 hours post cold exposure. n=6/group. **D)** Adipose tissue weight of control and Ad-*C3aR1* knockout female mice 6 hours post cold exposure. n=6/group. **E-F)** Hematoxylin and eosin staining (**E**) and UCP1 immunohistochemistry staining (**F**) of brown adipose tissue (BAT) sections from 10-weeek-old control and Ad-*C3aR1* knockout female mice at ambient temperature and following acute cold exposure. Images are shown at 20x magnification. Scale bar, 200 μm. Data are presented as mean± S.E.M. Unpaired two-tailed t test is used for comparison. *p < 0.05.

**Supplementary Table 1.**
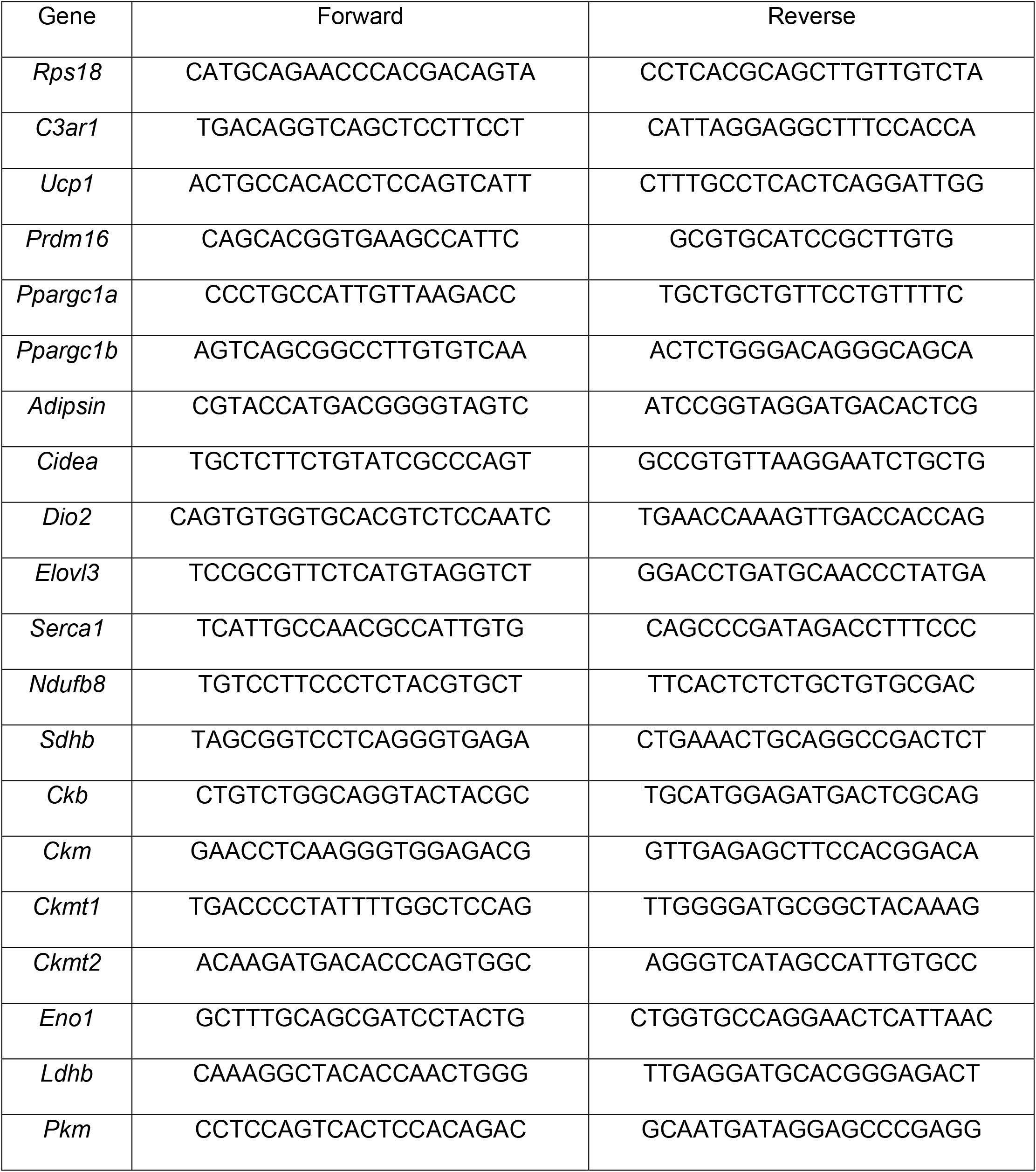
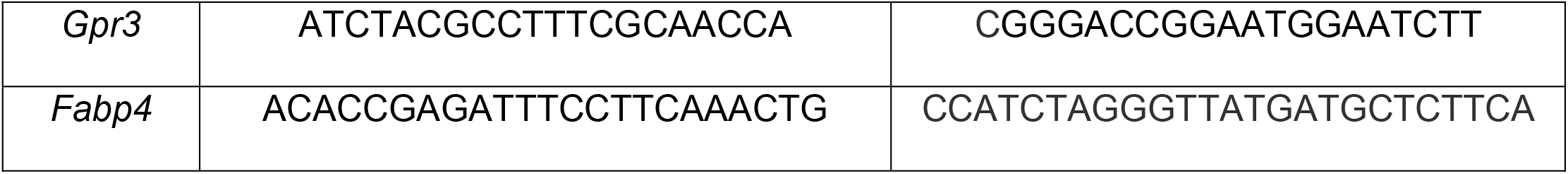
qPCR primers

## References

[1] T.M. Powell-Wiley, P. Poirier, L.E. Burke, J.P. Despres, P. Gordon-Larsen, C.J. Lavie, S.A. Lear, C.E. Ndumele, I.J. Neeland, P. Sanders, M.P. St-Onge, L. American Heart Association Council on, H. Cardiometabolic, C. Council on, N. Stroke, C. Council on Clinical, E. Council on, Prevention, C. Stroke, Obesity and Cardiovascular Disease: A Scientific Statement From the American Heart Association, Circulation 143(21) (2021) e984–e1010.

[2] P. Gonzalez-Muniesa, M.A. Martinez-Gonzalez, F.B. Hu, J.P. Despres, Y. Matsuzawa, R.J.F. Loos, L.A. Moreno, G.A. Bray, J.A. Martinez, Obesity, Nat Rev Dis Primers 3 (2017) 17034.

[3] S.Z. Yanovski, J.A. Yanovski, Long-term drug treatment for obesity: a systematic and clinical review, JAMA 311(1) (2014) 74–86.

[4] J. Rosenstock, C. Wysham, J.P. Frias, S. Kaneko, C.J. Lee, L. Fernandez Lando, H. Mao, X. Cui, C.A. Karanikas, V.T. Thieu, Efficacy and safety of a novel dual GIP and GLP-1 receptor agonist tirzepatide in patients with type 2 diabetes (SURPASS-1): a double-blind, randomised, phase 3 trial, Lancet 398(10295) (2021) 143–155.

[5] A.A. Rizvi, M. Rizzo, The Emerging Role of Dual GLP-1 and GIP Receptor Agonists in Glycemic Management and Cardiovascular Risk Reduction, Diabetes Metab Syndr Obes 15 (2022) 1023–1030.

[6] E.T. Chouchani, L. Kazak, B.M. Spiegelman, New Advances in Adaptive Thermogenesis: UCP1 and Beyond, Cell Metab 29(1) (2019) 27–37.

[7] B. Cannon, J. Nedergaard, Brown adipose tissue: function and physiological significance, Physiol Rev 84(1) (2004) 277–359.

[8] A.M. Cypess, A.P. White, C. Vernochet, T.J. Schulz, R. Xue, C.A. Sass, T.L. Huang, C. Roberts-Toler, L.S. Weiner, C. Sze, A.T. Chacko, L.N. Deschamps, L.M. Herder, N. Truchan, A.L. Glasgow, A.R. Holman, A. Gavrila, P.O. Hasselgren, M.A. Mori, M. Molla, Y.H. Tseng, Anatomical localization, gene expression profiling and functional characterization of adult human neck brown fat, Nat Med 19(5) (2013) 635–9.

[9] P. Cohen, S. Kajimura, The cellular and functional complexity of thermogenic fat, Nat Rev Mol Cell Biol 22(6) (2021) 393–409.

[10] J. Wu, P. Bostrom, L.M. Sparks, L. Ye, J.H. Choi, A.H. Giang, M. Khandekar, K.A. Virtanen, P. Nuutila, G. Schaart, K. Huang, H. Tu, W.D. van Marken Lichtenbelt, J. Hoeks, S. Enerback, P. Schrauwen, B.M. Spiegelman, Beige adipocytes are a distinct type of thermogenic fat cell in mouse and human, Cell 150(2) (2012) 366–76.

[11] Y.H. Lee, A.P. Petkova, A.A. Konkar, J.G. Granneman, Cellular origins of cold-induced brown adipocytes in adult mice, FASEB J 29(1) (2015) 286–99.

[12] J.Z. Long, K.J. Svensson, L. Tsai, X. Zeng, H.C. Roh, X. Kong, R.R. Rao, J. Lou, I. Lokurkar, W. Baur, J.J. Castellot, Jr., E.D. Rosen, B.M. Spiegelman, A smooth muscle-like origin for beige adipocytes, Cell Metab 19(5) (2014) 810–20.

[13] A.E. Pollard, D. Carling, Thermogenic adipocytes: lineage, function and therapeutic potential, Biochem J 477(11) (2020) 2071–2093.

[14] M. Shao, Q.A. Wang, A. Song, L. Vishvanath, N.C. Busbuso, P.E. Scherer, R.K. Gupta, Cellular Origins of Beige Fat Cells Revisited, Diabetes 68(10) (2019) 1874–1885.

[15] A. Sakers, M.K. De Siqueira, P. Seale, C.J. Villanueva, Adipose-tissue plasticity in health and disease, Cell 185(3) (2022) 419–446.

[16] A.M. Cypess, S. Lehman, G. Williams, I. Tal, D. Rodman, A.B. Goldfine, F.C. Kuo, E.L. Palmer, Y.H. Tseng, A. Doria, G.M. Kolodny, C.R. Kahn, Identification and importance of brown adipose tissue in adult humans, N Engl J Med 360(15) (2009) 1509–17.

[17] W.D. van Marken Lichtenbelt, J.W. Vanhommerig, N.M. Smulders, J.M. Drossaerts, G.J. Kemerink, N.D. Bouvy, P. Schrauwen, G.J. Teule, Cold-activated brown adipose tissue in healthy men, N Engl J Med 360(15) (2009) 1500–8.

[18] K.A. Virtanen, M.E. Lidell, J. Orava, M. Heglind, R. Westergren, T. Niemi, M. Taittonen, J. Laine, N.J. Savisto, S. Enerback, P. Nuutila, Functional brown adipose tissue in healthy adults, N Engl J Med 360(15) (2009) 1518–25.

[19] J.R. Dunkelberger, W.C. Song, Complement and its role in innate and adaptive immune responses, Cell Res 20(1) (2010) 34–50.

[20] Y. Mamane, C. Chung Chan, G. Lavallee, N. Morin, L.J. Xu, J. Huang, R. Gordon, W. Thomas, J. Lamb, E.E. Schadt, B.P. Kennedy, J.A. Mancini, The C3a anaphylatoxin receptor is a key mediator of insulin resistance and functions by modulating adipose tissue macrophage infiltration and activation, Diabetes 58(9) (2009) 2006–17.

[21] J.C. Lo, S. Ljubicic, B. Leibiger, M. Kern, I.B. Leibiger, T. Moede, M.E. Kelly, D. Chatterjee Bhowmick, I. Murano, P. Cohen, A.S. Banks, M.J. Khandekar, A. Dietrich, J.S. Flier, S. Cinti, M. Bluher, N.N. Danial, P.O. Berggren, B.M. Spiegelman, Adipsin is an adipokine that improves beta cell function in diabetes, Cell 158(1) (2014) 41–53.

[22] K. Shim, R. Begum, C. Yang, H. Wang, Complement activation in obesity, insulin resistance, and type 2 diabetes mellitus, World J Diabetes 11(1) (2020) 1–12.

[23] L.N. Choy, B.S. Rosen, B.M. Spiegelman, Adipsin and an endogenous pathway of complement from adipose cells, J Biol Chem 267(18) (1992) 12736–41.

[24] N. Gomez-Banoy, J.S. Guseh, G. Li, A. Rubio-Navarro, T. Chen, B. Poirier, G. Putzel, C. Rosselot, M.A. Pabon, J.P. Camporez, V. Bhambhani, S.J. Hwang, C. Yao, R.J. Perry, S. Mukherjee, M.G. Larson, D. Levy, L.E. Dow, G.I. Shulman, N. Dephoure, A. Garcia-Ocana, M. Hao, B.M. Spiegelman, J.E. Ho, J.C. Lo, Adipsin preserves beta cells in diabetic mice and associates with protection from type 2 diabetes in humans, Nat Med 25(11) (2019) 1739–1747.

[25] J. Lim, A. Iyer, J.Y. Suen, V. Seow, R.C. Reid, L. Brown, D.P. Fairlie, C5aR and C3aR antagonists each inhibit diet-induced obesity, metabolic dysfunction, and adipocyte and macrophage signaling, FASEB J 27(2) (2013) 822–31.

[26] J. Wu, P. Cohen, B.M. Spiegelman, Adaptive thermogenesis in adipocytes: Is beige the new brown?, Gene Dev 27(3) (2013) 234–250.

[27] L.A. Trouw, M.C. Pickering, A.M. Blom, The complement system as a potential therapeutic target in rheumatic disease, Nat Rev Rheumatol 13(9) (2017).

[28] S.A. Zwarthoff, E.T.M. Berends, S. Mol, M. Ruyken, P.C. Aerts, M. Jozsi, C.J.C. de Haas, S.H.M. Rooijakkers, R.D. Gorham, Functional Characterization of Alternative and Classical Pathway C3/C5 Convertase Activity and Inhibition Using Purified Models, Frontiers in Immunology 9 (2018).

[29] K.R. Mayilyan, Complement genetics, deficiencies, and disease associations, Protein Cell 3(7) (2012) 487–96.

[30] M. Gaya da Costa, F. Poppelaars, C. van Kooten, T.E. Mollnes, F. Tedesco, R. Wurzner, L.A. Trouw, L. Truedsson, M.R. Daha, A. Roos, M.A. Seelen, Age and Sex-Associated Changes of Complement Activity and Complement Levels in a Healthy Caucasian Population, Front Immunol 9 (2018) 2664.

[31] N. Kamitaki, A. Sekar, R.E. Handsaker, H. de Rivera, K. Tooley, D.L. Morris, K.E. Taylor, C.W. Whelan, P. Tombleson, L.M.O. Loohuis C. Schizophrenia Working Group of the Psychiatric Genomics, M. Boehnke, R.P. Kimberly, K.M. Kaufman, J.B. Harley, C.D. Langefeld, C.E. Seidman, M.T. Pato, C.N. Pato, R.A. Ophoff, R.R. Graham, L.A. Criswell, T.J. Vyse, S.A. McCarroll, Complement genes contribute sex-biased vulnerability in diverse disorders, Nature 582(7813) (2020) 577–581.

[32] A. Pfeifer, L.S. Hoffmann, Brown, beige, and white: the new color code of fat and its pharmacological implications, Annu Rev Pharmacol Toxicol 55 (2015) 207–27.

[33] F. Forneris, D. Ricklin, J. Wu, A. Tzekou, R.S. Wallace, J.D. Lambris, P. Gros, Structures of C3b in complex with factors B and D give insight into complement convertase formation, Science 330(6012) (2010) 1816–20.

[34] W.D. Campbell, E. Lazoura, N. Okada, H. Okada, Inactivation of C3a and C5a octapeptides by carboxypeptidase R and carboxypeptidase N, Microbiol Immunol 46(2) (2002) 131–4.

[35] A.B. Molofsky, J.C. Nussbaum, H.E. Liang, S.J. Van Dyken, L.E. Cheng, A. Mohapatra, A. Chawla, R.M. Locksley, Innate lymphoid type 2 cells sustain visceral adipose tissue eosinophils and alternatively activated macrophages, J Exp Med 210(3) (2013) 535–49.

[36] D. Wu, A.B. Molofsky, H.E. Liang, R.R. Ricardo-Gonzalez, H.A. Jouihan, J.K. Bando, A. Chawla, R.M. Locksley, Eosinophils sustain adipose alternatively activated macrophages associated with glucose homeostasis, Science 332(6026) (2011) 243–7.

[37] R.F. Ritchie, G.E. Palomaki, L.M. Neveux, O. Navolotskaia, T.B. Ledue, W.Y. Craig, Reference distributions for complement proteins C3 and C4: a practical, simple and clinically relevant approach in a large cohort, J Clin Lab Anal 18(1) (2004) 1–8.

[38] A.S. Silva, A.G. Teixeira, L. Bavia, F. Lin, R. Velletri, R. Belfort, Jr., L. Isaac, Plasma levels of complement proteins from the alternative pathway in patients with age-related macular degeneration are independent of Complement Factor H Tyr(4)(0)(2)His polymorphism, Mol Vis 18 (2012) 2288–99.

[39] E. Filatov, L.I. Short, M.A.M. Forster, S.S. Harris, E.N. Schien, M.C. Hughes, D.L. Cline, C.J. Appleby, S.L. Gray, Contribution of thermogenic mechanisms by male and female mice lacking pituitary adenylate cyclase-activating polypeptide in response to cold acclimation, Am J Physiol Endocrinol Metab 320(3) (2021) E475–E487.

[40] A.P. Frank, B.F. Palmer, D.J. Clegg, Do estrogens enhance activation of brown and beiging of adipose tissues?, Physiol Behav 187 (2018) 24–31.

[41] K. Kaikaew, A. Grefhorst, J.A. Visser, Sex Differences in Brown Adipose Tissue Function: Sex Hormones, Glucocorticoids, and Their Crosstalk, Front Endocrinol (Lausanne) 12 (2021) 652444.

[42] V. Benz, M. Bloch, S. Wardat, C. Bohm, L. Maurer, S. Mahmoodzadeh, P. Wiedmer, J. Spranger, A. Foryst-Ludwig, U. Kintscher, Sexual dimorphic regulation of body weight dynamics and adipose tissue lipolysis, PLoS One 7(5) (2012) e37794.

[43] A. Cumpelik, D. Heja, Y. Hu, G. Varano, F. Ordikhani, M.P. Roberto, Z. He, D. Homann, S.A. Lira, D. Dominguez-Sola, P.S. Heeger, Dynamic regulation of B cell complement signaling is integral to germinal center responses, Nat Immunol 22(6) (2021) 757–768.

